# Extensive Diversity of RNA Viruses in Australian Ticks

**DOI:** 10.1101/386573

**Authors:** Erin Harvey, Karrie Rose, John-Sebastian Edena, Nathan Lo, Thilanka Abeyasuriya, Mang Shi, Stephen L. Doggett, Edward C. Holmes

**Affiliations:** Marie Bashir Institute for Infectious Diseases and Biosecurity, Charles Perkins Centre, School of Life and Environmental Sciences and Sydney Medical School, The University of Sydney, Sydney, NSW 2006, Australia.; Australian Registry of Wildlife Health, Taronga Conservation Society Australia, Mosman, NSW 2088, Australia.; Centre for Virus Research, Westmead Institute for Medical Research, Westmead NSW 2145.; School of Life and Environmental Sciences and Sydney Medical School, The University of Sydney, Sydney, NSW 2006, Australia.; Department of Medical Entomology, NSWHP-ICPMR, Westmead Hospital, Westmead, NSW 2145, Australia.

**Keywords:** ticks, *Ixodes holocyclus*, RNA virus, virome, phylogeny, marsupial, coltivirus

## Abstract

Understanding the microbiome of ticks in Australia is of considerable interest given the ongoing debate over whether Lyme disease, and its causative agent the bacterium *Borrelia burgdorferi* senso lato, are present in Australia. The diversity of bacteria infecting Australian ticks has been the subject of a number of studies using both culture and metagenomics based techniques. However, little is known about the virome of Australian ticks, including those that may have the potential to infect mammalian species. We used a meta-transcriptomics approach to reveal the viral diversity within Australian ticks collected from two locations on the central-east coast of Australia, including metropolitan Sydney. From this we identified 19 novel species of RNA virus belonging to 10 families, as well as one previously described RNA virus. The majority of these viruses clustered phylogenetically with arthropod-associated viruses suggesting that they do not utilize mammalian hosts. However, two novel viruses discovered in ticks feeding on bandicoot marsupials clustered closely within the mammalian associated *Hepacivirus* and *Pestivirus* genera (*Flaviviridae*). Notably, another bandicoot tick yielded a novel *Coltivirus* (*Reoviridae*) – a group of largely tick-associated viruses containing the known human pathogen Colorado tick fever virus and its relative Eyach virus. Importantly, our transcriptomic data provided no evidence for the presence of *B. burgdorferi* s.l.. in any tick sample, providing further evidence against the presence of Lyme Disease in Australia. In sum, this study reveals that Australian ticks harbor a diverse virome, including some viruses that merit additional screening in the context of emerging infectious disease.

**IMPORTANCE:** Each year a growing number of individuals along the east coast of Australia experience debilitating disease following tick bites. As there is no evidence for the presence of the causative agent of Lyme disease, *Borrelia Burgdorferi* seno lato, in Australian ticks, the etiological basis of this disease syndrome remains controversial. To characterize the viruses associated with Australian ticks, particularly those that might be associated with mammalian infection, we performed unbiased RNA sequencing on 146 ticks collected across two locations along the coast of New South Wales, Australia. This revealed 19 novel RNA viruses from a diverse set of families. Notably, three of these viruses were related to known mammalian viruses, including one that fell within the genus *Coltivirus* and related to the human pathogen, Colorado tick fever virus.

---

Ticks are one of the most important vectors of infectious disease in humans, wildlife, and domestic animals (1), in part due to their ability to harbor multiple viruses, bacteria and eukaryotic parasites, often simultaneously (2). Due to ecological changes affecting the life cycle and range of ticks, such as increased encroachment by humans on tick habitats and the impact of climate change in reducing tick mortality in winter and extending the active period, ticks are now an increasingly important vector of zoonotic disease globally (3). For example, cases of tick-borne bacterial infection, particularly Lyme disease, are becoming increasingly frequent across parts of North America and Europe (4, 5). In Australia, Lyme disease is not recognized as endemic by the scientific nor medical communities, although controversy over its presence along the eastern coast of Australia has raged since the mid-1980s (6-8). Importantly, the species of ticks known to act as a vector of the disease are not found in Australia, and detailed studies involving microscopy, culturing, PCR, and metagenomic techniques indicate that the causative bacterial agent, *Borrelia burgdorferi* senso lato, is not present in Australian ticks (6, 7). In addition, a North American strain of *B. burgdorferi* could not be transmitted by the Australian paralysis tick, *Ixodes holocyclus*, following experimental infection (9). Finally, and of most note, *B. burgdorferi* has not been isolated from patients suffering the physical symptoms of disease following bites from Australian ticks (10). Hence, the cause of many human tick-borne disease events in Australia remains both unknown and controversial.

A number of studies have used bacterial 16S rRNA-based metagenomics, as well as PCR and immunoassay-based methods, to search for known or novel tick-associated bacterial pathogens that may be present in Australian ticks (11-13). Additional studies have used more powerful, bulk RNA sequencing-based metagenomic approaches to identify viruses in diverse tick populations, although not those from Australia (14). Although ticks carry a diverse virome and are associated with the transmission of viral pathogens in parts of Europe, North America, Asia, and Africa (15-19), there has been no comprehensive metagenomic study of the virome of ticks in Australia.

Australia and its wildlife have largely evolved in isolation for over 46 million years and are characteristically unique and diverse. Hence, a survey of tick viruses in the context of tick-borne disease might be expected to identify equally diverse and unexpected organisms. (20). Indeed, a number of novel tick borne-viruses, such as Albatross Island virus and Saumarez Reef virus, have been identified using culture-dependent methods in Australian wildlife (21, 22). However, these methods are inherently limited by the number of viruses that can be cultured, and have often focused on microbial isolation following die-off events in the ticks host population. The description of such a small number of tick associated viruses means that there is a paucity of knowledge about the natural virome of ticks in Australia.

We have recently shown that meta-transcriptomics (i.e. bulk shotgun RNA-sequencing) is a powerful tool for the discovery of novel viruses and other microbial pathogens from both host and tick-derived samples (14, 23-25). Importantly, because total RNA is sequenced, meta-transcriptomics provides information on the total “infectome” of the sample and can be used to the identify viruses, bacteria, fungi, and other eukaryotes present within a sample (26). The method also provides quantitative data on virus abundance that may be used to identify potential pathogens (26). To better understand the natural virome of Australian ticks and their evolutionary relationships, with a specific focus on RNA viruses, we performed unbiased RNA sequencing on 146 ticks from three tick species collected from native lizards, marsupials, and rodents, from introduced rodents, as well as from unfed nymphal ticks. In particular, we sought to determine whether Australian ticks harbor any viruses from families previously associated with human and other mammalian disease. All tick specimens were collected from the North Shore region of Sydney and the South Coast region of New South Wales (NSW), and the data obtained provide important insights into the remarkable diversity of tick-associated RNA viruses.

## RESULTS

### Confirmation of tick species

In total, 146 ticks were collected across two locations in New South Wales, Australia, during 2016 and 2017. The samples represented three tick species common to the east coast of Australia – *I. holocyclus, I. trichosuri* and *Amblyomma moreliae* (Table 1). Ticks were pooled into 11 libraries based on tick species, collection location, host species, and life-cycle stage where possible (Table 1). RNA-sequencing generated between 32,280,822 and 59,377,716 reads per library which were assembled *de novo* into between 159,626 and 505,268 contigs per library (Table 1). Tick mitochondrial COX1 sequences were identified from the assembled contigs, and used to confirm the species identification of ticks in each library (Fig. 1).

**Table 1.**
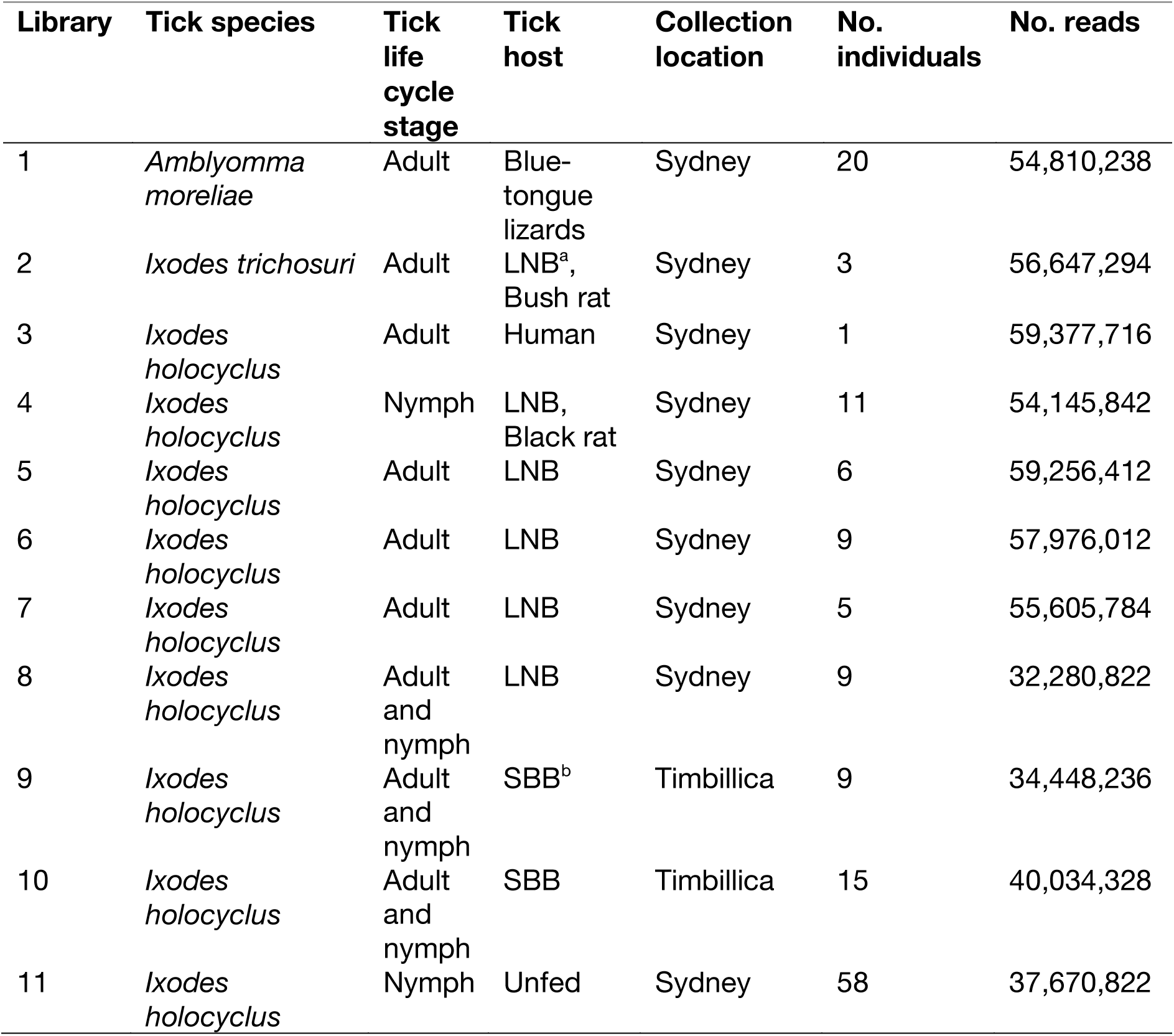
Summary details of each RNA sequencing library.

**FIG 1.**
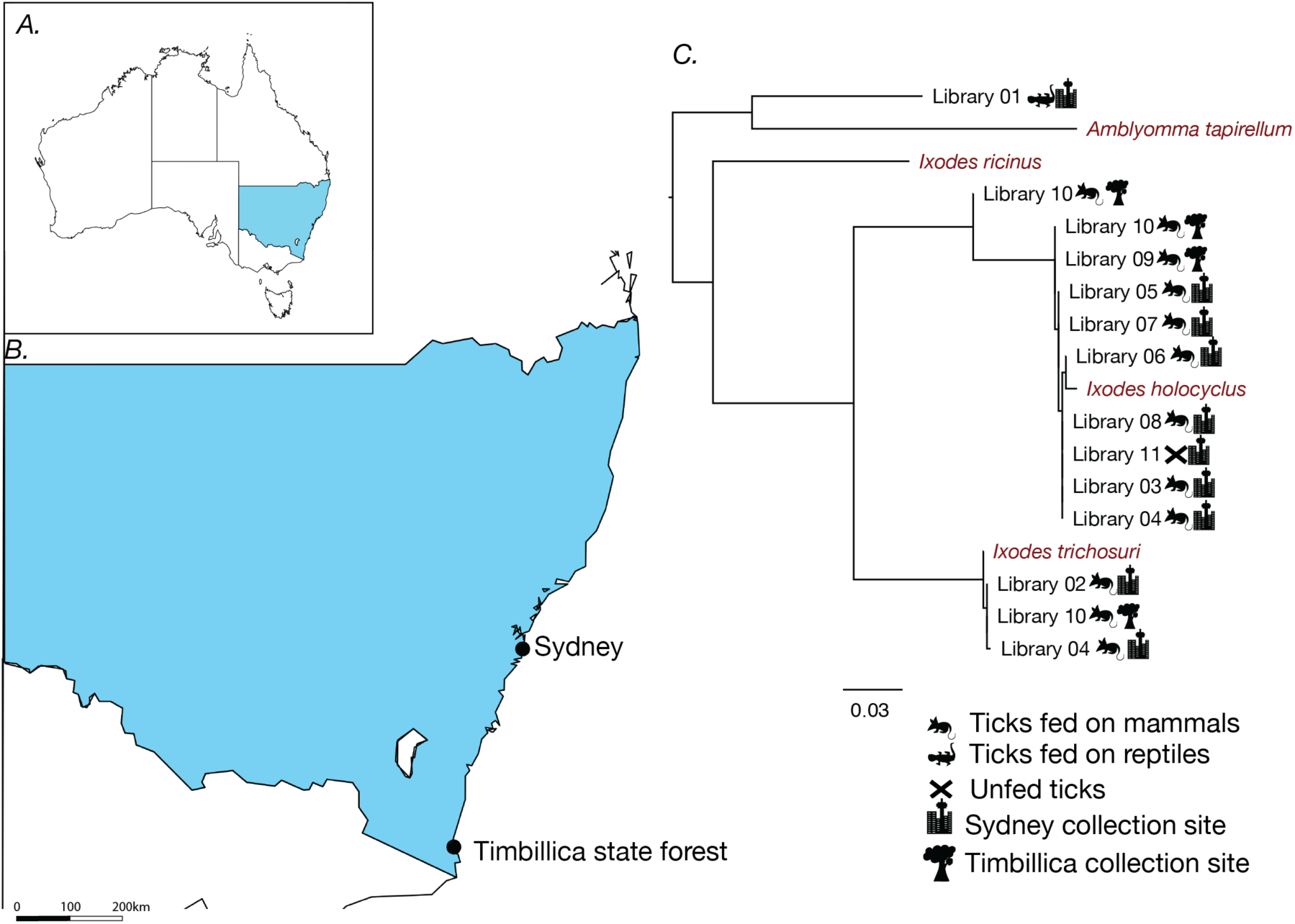
A. Map of Australia showing the location of New South Wales, where sampling took place. B. Locations of tick sampling sites marked by solid black dots on a map of New South Wales, Australia. C. Maximum likelihood phylogeny of the mitochondrial cytochrome c oxidase subunit I (COX 1) gene from each pooled library, with the sampling species (and unfed ticks) and location indicated. Reference COX1 sequences from a variety of tick species are shown in red. The scale bar indicates number of nucleotide substitutions per site. The tree was mid-pointed rooted for clarity only.

A single COX1 sequence was identified for libraries 3, 5-9 and 11, all of which were all closely related to the *I. holocyclus* COX1 reference sequence taken from NCBI (Fig. 1). In contrast, libraries 4 and 10 contained multiple distinct COX1 sequences suggesting that there may have been more than one species of tick included in the library that had been incorrectly classified. No reference sequence was available for the tick species *A. moreliae*. Accordingly, the most closely related species represented in the NCBI refseq database, *Amblyomma tapirellum,* was included in the phylogeny as a proxy, although it is important to note that this species of tick is not found in Australia. Importantly, none of the COX1 sequences identified in the 11 transcriptomes showed strong sequence similarity to *Ixodes ricinus*, the main tick species that acts as a vector of *B. burgdorferi* s.l. in western Europe, nor to *Ixodes scapularius*, *Ixodes persulcatus* or *Ixodes pacificus*.

### The tick virome

Overall, 16 complete viral genomes were identified within the tick data set collected here using a series of BLAST searches based on protein sequence similarity. An additional four partial viral genomes were also identified. Of these 20 complete or partial genomes, 19 contained previously undescribed viruses, although all fell within known viral families or orders, specifically: *Mononegavirales, Orthomyxoviridae, Picornaviridae, Flaviviridae, Narnaviridae, Luteoviridae, Virgaviridae, Reoviridae, Phenuiviridae*, and *Partitiviridae*. All viruses identified across the 11 libraries were RNA viruses with no DNA intermediate, but the full diversity of RNA virus genome structures was represented; that is, positive-sense single-stranded RNA (+ssRNA), negative-sense single-stranded RNA (-ssRNA), and double-stranded RNA (dsRNA), segmented, and non-segmented. It is unlikely that any of these viruses are endogenous viral elements present within the tick genome as they were identified by ORFs unbroken by stop codons and none showed nucleotide sequence similarity to tick gene sequences within the NCBI BLAST database.

The housekeeping gene COX1 was used to measure the abundance of the invertebrate (i.e. likely host) transcripts in each library. Accordingly, COX1 abundance ranged from 0.15% to 1.1% of the total number of reads (from which rRNA was already excluded; Fig. 2). We also calculated the proportion of virus reads as a fraction of total reads. This revealed that while the number of viral reads in each library ranged from 47 to 268,775 (0.00017% to 0.29% of the number of non-ribosomal reads within a library), it was not related to the abundance of host genes (Fig. 2). However, the number of virus species in each library, which varied from one to six, appears to be associated with the percentage abundance of virus reads (Fig. 2).

**FIG 2.**
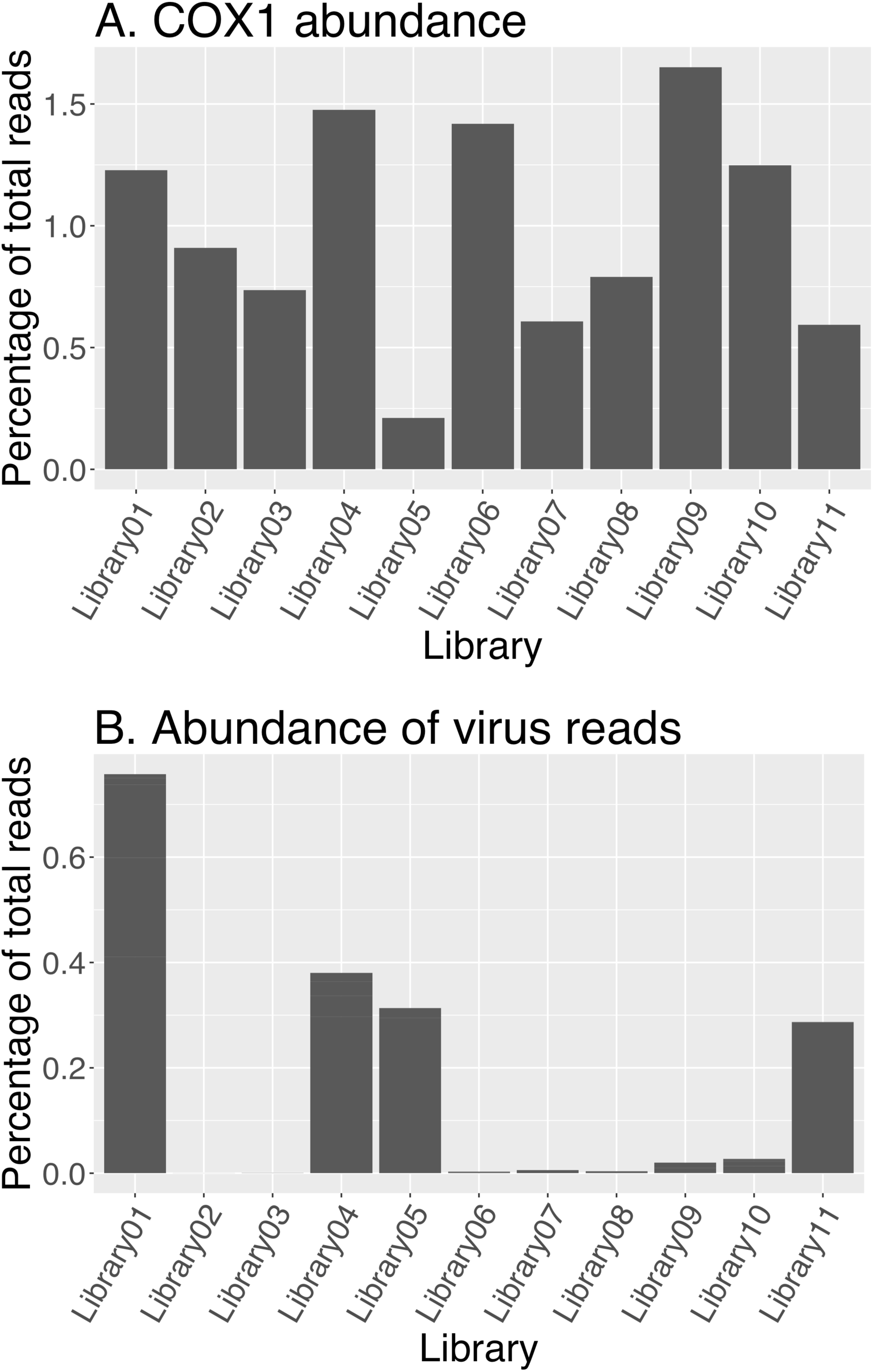
A. Percentage of reads aligning to the tick mitochondrial COX1 reference sequence. B. Percentage of reads aligning to RNA virus genomes in each library. Library composition is described in Table 1.

Viruses from the unclassified *Chuvirus* group (-ssRNA) and the *Picornaviridae* (+ssRNA) were the most abundant, while viruses from the *Flaviviridae* (+ssRNA) were the least abundant. In libraries containing multiple viruses, the abundance of each species varied substantially. For example, in Library 1, Sydney tick mononega-like virus 1 represented 54% of the total number of virus reads, while Sydney tick mononega-like virus 4 represented only 0.9% of the total virus reads in that library (Fig. 3).

**FIG 3.**
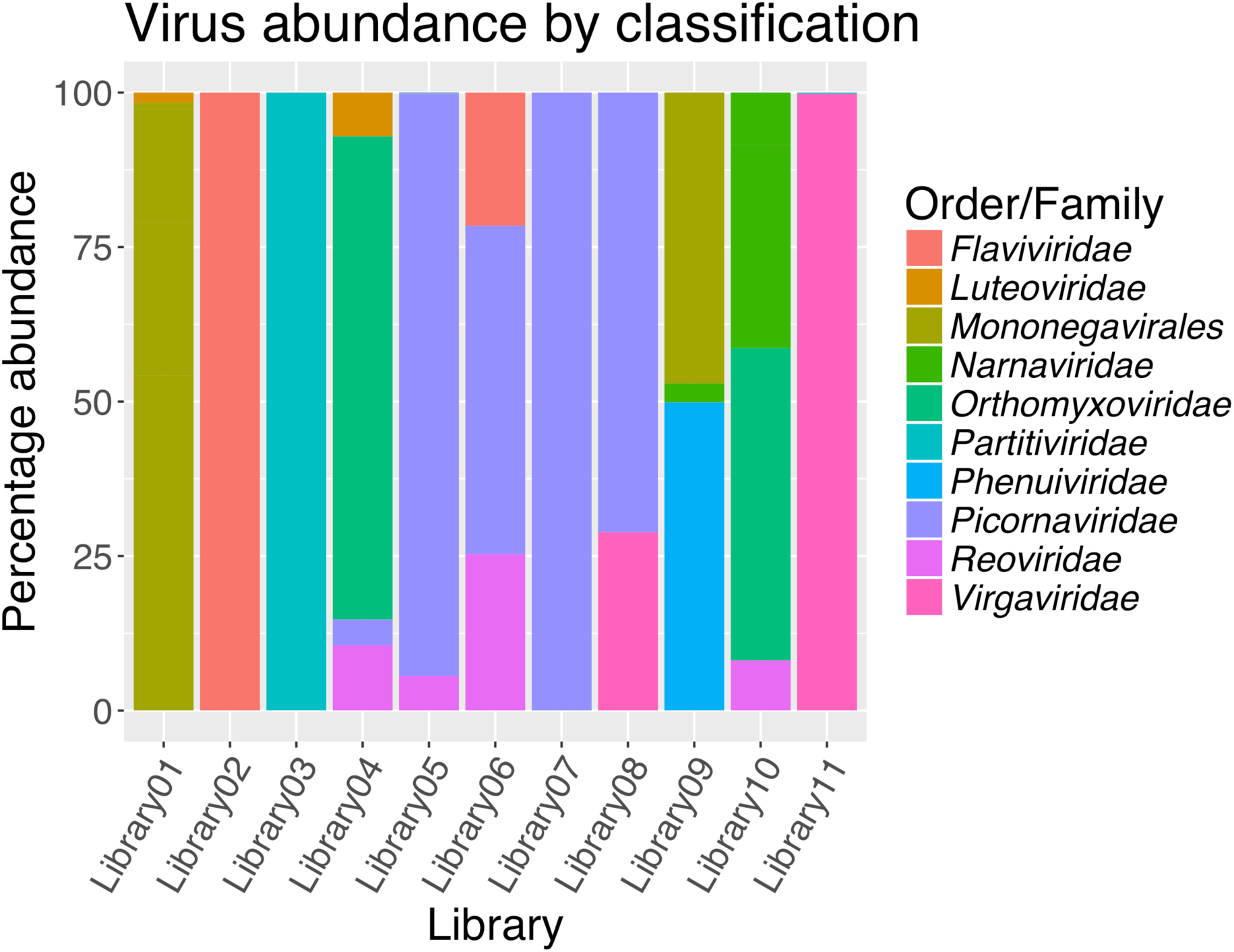
The abundance of each virus family or order in each sequencing library. Abundance is shown as the percentage of the total number of viral reads in each library.

The evolutionary history of each novel virus was investigated using phylogenetic analysis. In particular, this analysis was used to infer further information about the likely host of each virus as well the relationship of each virus to previously characterized viruses. The 19 novel viruses described here showed varying degrees of similarity to those previously identified.

### Positive-sense RNA viruses

Our data contained genomic evidence for the presence of nine +ssRNA viruses belonging to five virus families, representing just under half of the viruses identified in this study. Importantly, this includes two flavi-like viruses potentially associated with mammalian hosts, rather than the tick itself, based on their position within the *Flaviviridae* phylogeny (Fig. 4) and their low abundance (Fig. 2). Specifically, both Sydney hepaci-like virus 1 and Sydney pesti-like virus 1 exhibited extremely low abundance in the data set and only partial viral genomes were identified (Table 2). Despite this, enough of the viral genome was assembled to perform robust phylogenetic analysis, which revealed that these viruses were closely related to rodent hepacivirus and Norway rat pestivirus, respectively (Fig. 4). Their relatively close evolutionary relationship to other rodent associated viruses gives additional support to the idea that these viruses are associated with the blood meal contained within the engorged ticks and may not be infecting (i.e. replicating in) the ticks themselves. Notably, Sydney pesti-like virus 1 was identified in *I. trichosuri* ticks that had fed on long-nosed bandicoots (a marsupial) and native bush rats (Table 1). Similarly, Sydney hepaci-like virus 1 was found in *I. holocyclus* collected from long-nosed bandicoots (Table 1).

**Table 2.** Name and description of the novel RNA viruses found in this study.

| **Virus name** | **Virus family/**<br>**order** | **Genome**<br>**length /**<br>**contig**<br>**length** | **No.**<br>**segments** | **Present in**<br>**libraries (L)** |
| --- | --- | --- | --- | --- |
| Sydney tick mononega-like virus 1 | *Mononegavirales* | 9835 | 1 | L1 |
| Sydney tick mononega-like virus 2 | *Mononegavirales* | 9047 | 1 | L1 |
| Sydney tick mononega-like virus 3 | *Mononegavirales* | 11192 | 1 | L1 |
| Sydney tick luteo-like virus 1 | *Luteoviridae* | 2632 | 1 | L1 |
| Sydney tick mononega-like virus 4 | *Mononegaviridae* | 8667 | 1 | L1 |
| Sydney tick partiti-like virus 1 | *Partitiviridae* | 1770 | 1 | L3, L5, L11 |
| Sydney tick luteo-like virus 2 | *Luteoviridae* | 4373 | 2 | L4 |
| Sydney tick colti-like virus 1 | *Reoviridae* | 26411 | 11 | L5, L6 |
| Sydney tick picorna-like virus 1 | *Picornaviridae* | 5202<sup>a</sup> | 1 | L5 |
| Sydney tick virga-like virus 1 | *Virgaviridae* | 11176 | 1 | L8, L11 |
| Sydney hepaci-like virus 1 | *Flaviviridae* | 1343<sup>a</sup> | 1 | L5, L6 |
| Sydney pesti-like virus 1 | *Flaviviridae* | 2194<sup>a</sup> | 1 | L2 |
| Timbillica tick Phenui-like virus 1 | *Phenuiviridae* | 8909 | 2 | L9 |
| Timbillica tick Mononega-like virus 1 | *Mononegavirales* | 11018 | 1 | L9 |
| Timbillica tick narna-like virus 1 | *Narnaviridae* | 1733<sup>a</sup> | 1 | L9 |
| Timbillica tick narna-like virus 2 | *Narnaviridae* | 3248 | 1 | L10 |
| Timbillica tick narna-like virus 3 | *Narnaviridae* | 2406 | 1 | L10 |
| Sydney tick orthomyxo-like virus 1 | *Orthomyxoviridae* | 10498 | 5 | L4, L10 |
| Sydney tick reo-like virus 1 | *Reoviridae* | 13739 | 6 | L4, L10 |

**FIG 4.**
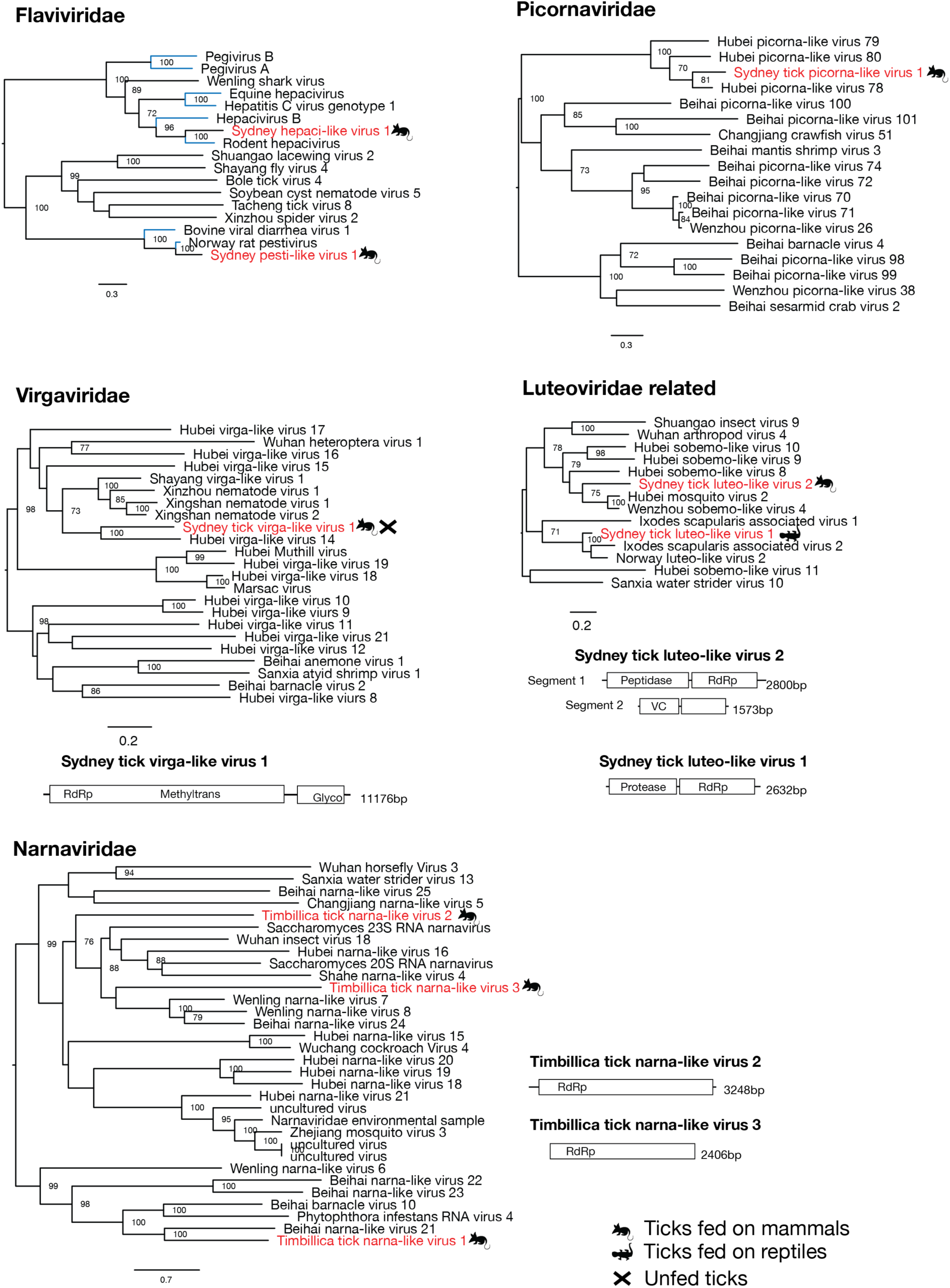
Phylogenetic relationships and genomic structure of the +ssRNA viruses sampled in this study. All trees are scaled according to the number of amino acid substitutions per site. The trees are midpoint rooted for clarity only and bootstrap values (>70%) are shown. Viruses determined here are shown in red. Animal symbols denote the vertebrate host the tick was taken from or whether the virus was sampled from an unfed tick. Virus lineages previously shown to infect mammals or mammalian cells are shown in blue. Genome diagrams provide information on the length of each genomic segment, the number of ORFs and predicted conserved protein structures. For viruses where no full genome was identified in the data, no such diagram is provided.

In contrast, all the remaining +ssRNA viruses identified were closely related to arthropod-associated viruses. Sydney tick virga-like virus 1 fell within the virga-like virus cluster and was most closely related to a virus found in spiders from China (Figure 4) (23). The genome of Sydney tick virga-like virus 1 is 11,176 nt long and encodes two predicted ORFs. This virus represented 99.8% of viral reads and 0.2% of the total reads in library 11, and was also present in low abundance in library 8 (Table 2, Fig. 2). Similarly, Sydney tick luteo-like virus 2 and Sydney tick luteo-like virus 1 clustered with viruses of the *Luteoviridae* family. While Sydney tick luteo-like virus 2, a bi-segmented virus with 4 ORFs, is most closely related to mosquito-associated viruses, Sydney tick luteo-like virus 1 appears to comprise a single segment with two ORFs that groups closely with other *Ixodes* associated viruses (Fig. 4). While Sydney tick luteo-like virus 2 was associated with *I. holocyclus* collected from long-nosed bandicoots, Sydney tick luteo-like virus 1 was identified in lizard ticks, and was the only virus found in the *Amblyomma moreliae* pool. Despite the divergent phylogenetic position of *A. moreliae*, Sydney tick luteo-like virus 1 was closely related to viruses associated with *Ixodes* ticks (Fig. 4). Finally, Sydney tick picorna-like virus 1 groups with other Picorna-like viruses identified in Chinese arthropods (Fig. 4) (23). A full genome of this virus was not recovered from the data, likely due its low abundance, as the 5202 bp fragments represented only 0.14% of viral reads in the library.

Viruses showing similarity to the Narna-like virus family were only identified in ticks collected in southern NSW (Timbillica), and all three species were extremely divergent from those viruses described previously. Amino acid sequence similarity to the most closely related virus species ranged between 24 and 32% for these three viruses across the RdRp region. These viruses comprised a single genome segment with one predicted ORF containing an RdRp-like region, typical of viruses within this family which have no known structural proteins (27). Complete genomes were recovered for two of these viruses, designated Timbillica tick narna-like virus 2 and Timbillica tick narna-like virus 3, while only fragments of Timbillica tick narna-like virus 1 were recovered from this data set.

The previously identified *I. holocyclus* iflavirus (28), found in *I. holocyclus* using a viral RNA isolation method, was present in five libraries (libraries 4-8). This is the only previously described virus found in any of the 11 data sets generated here. This virus was only identified in *I. holocyclus* collected in the Sydney region.

### Double-strand RNA viruses

Our data contained genomic evidence for the presence of three dsRNA viruses from two families. Sydney tick partiti-like virus 1 clustered with uncharacterized partiti-like viruses isolated from arthropods. On the phylogeny this group of viruses fell between deltapartitiviruses and gammapartitiviruses which are known to infect plants and fungi, respectively (Fig. 5). For this reason, and because partitiviruses are known to infect a wide range of hosts from protozoans to fungi (29), it was impossible to determine the likely host of these viruses. A single segment with a single ORF showed only approximately 50% amino acid similarity over the conserved RdRp region. This virus was found in three libraries of *I. holocyclus* ticks from the Sydney region (Table 1).

**FIG 5.**
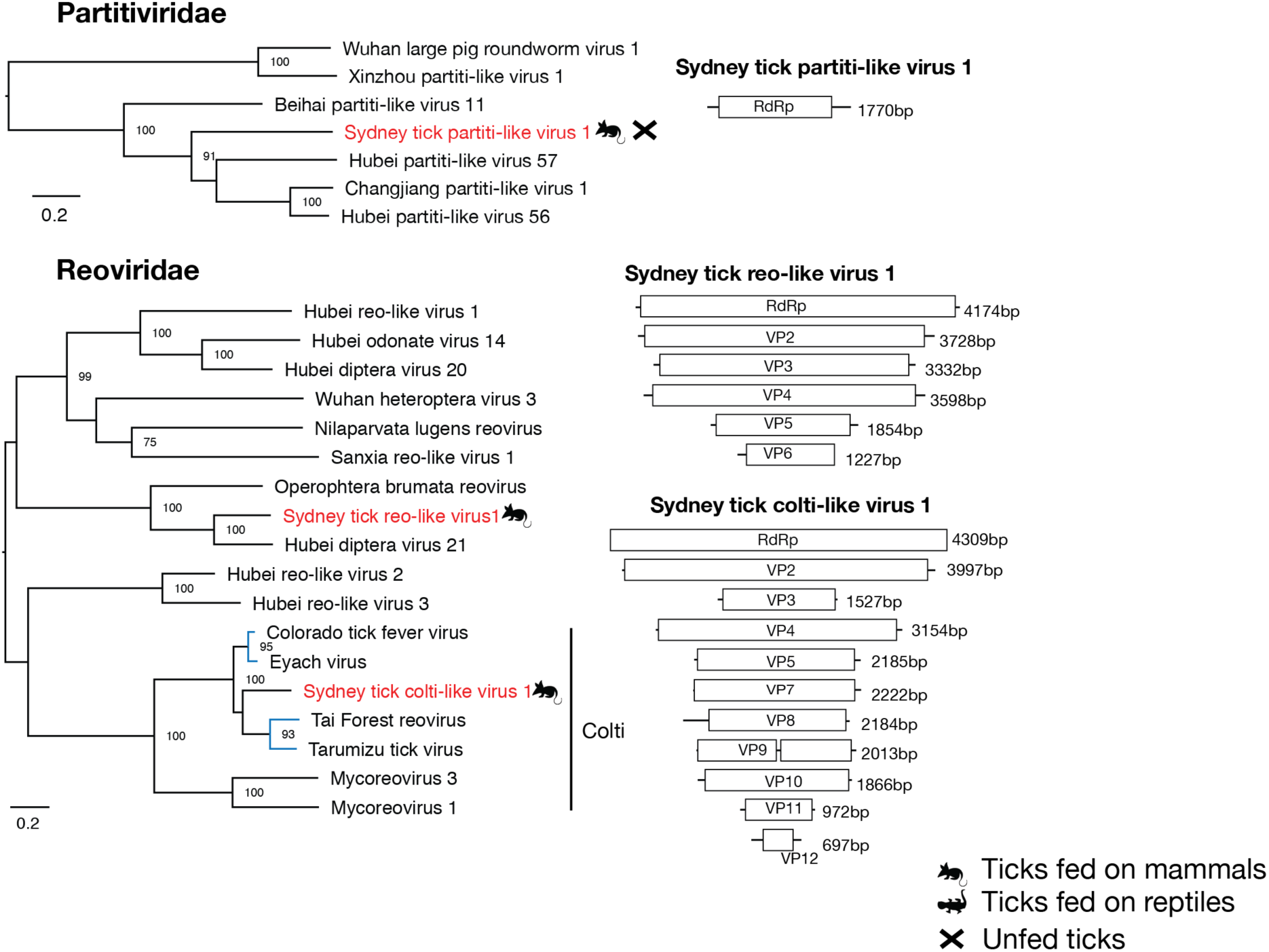
Phylogenetic relationships and genomic structure of the dsRNA viruses sampled in this study. All trees are scaled according to the number of amino acid substitutions per site. The trees were midpoint rooted for clarity only and bootstrap values (>70%) are shown. Viruses determined here are shown in red. Animal symbols denote the vertebrate host the tick was taken from or whether the virus was sampled from an unfed tick. Virus lineages previously shown to infect mammals or mammalian cells are shown in blue. Genome diagrams provide information on the length of each genomic segment, the number of predicted ORFs and predicted conserved protein structures.

Sydney tick reo-like virus 1 and Sydney tick colti-like virus 1 clustered with the *Reoviridae*, a family of segmented RNA viruses (Fig. 5). Sydney tick reo-like virus 1 was found to have six segments encoding six ORFs and was most closely related to other reoviruses associated with flies and moths (23). This is one of two viruses found in *I. holocyclus* collected from both locations. Of greater interest was Sydney tick colti-like virus 1 that grouped with the subfamily of tick-associated *Coltiviruses*. Importantly, two coltiviruses have been implicated in human disease in North America and Europe – Colorado tick fever virus (CTFV) and Eyach virus (15, 16). The other two members of the subfamily were recently identified in viral culture isolated from ticks in Japan and the blood of free-tailed bats in Côte d’lvoire (30, 31). In total, 11 segments of Sydney tick colti-like virus 1 were identified containing 12 ORFs, with segment 9 containing two predicted ORFs separated by a single stop codon. It is currently unclear if there is a 12^th^ segment containing a viral protein similar to that of viral protein 6 in CTFV that cannot be identified through amino acid similarity, or if this segment is not present in this virus. It has been previously observed that segment 6 of the coltiviruses does not exhibit amino acid sequence similarity to other viruses within the genus (30).

### Negative-sense RNA viruses

The full genomes of seven -ssRNA viruses were identified within four of the 11 libraries produced in this study. Two of these viruses fell within the *Phenuiviridae* and *Orthomyxoviridae*, while the remaining five fell within the order *Mononegavirales* and clustered with the unclassified mononegavirales-chuvirus group (32). Timbillica tick Phenui-like virus 1 clustered within the *Phenuiviridae*, but is extremely divergent from previously described viruses in this family, with only 36% amino acid similarity to the most closely related virus (Blacklegged tick phlebovirus 3). The most closely related viruses are also tick-associated and were isolated from ticks in Norway and North America (Fig. 6) (14, 33). Timbillica tick Phenui-like virus 1 grouped more closely with viruses found to infect ticks of the *Ixodes* genus rather than with Albatross Island virus (previously known as Hunter Island virus) identified in avian ticks collected from albatross in Tasmania (21). Only two segments with a single ORF each were identified (Fig. 6). This could be due to either an absence or lack of identification through amino acid similarity searches of the third segment typically associated with phleboviruses. Sydney tick orthomyxo-like virus 1 was the only virus identified in both the Timbillica and Sydney ticks, although the virus was far more abundant in the library containing ticks from the Sydney region (Fig. 3). Interestingly, this virus was most closely related to other arthropod-associated viruses, and contained at least five segments, each with a single ORF, as was typical of other related viruses (Fig. 6).

**FIG 6.**
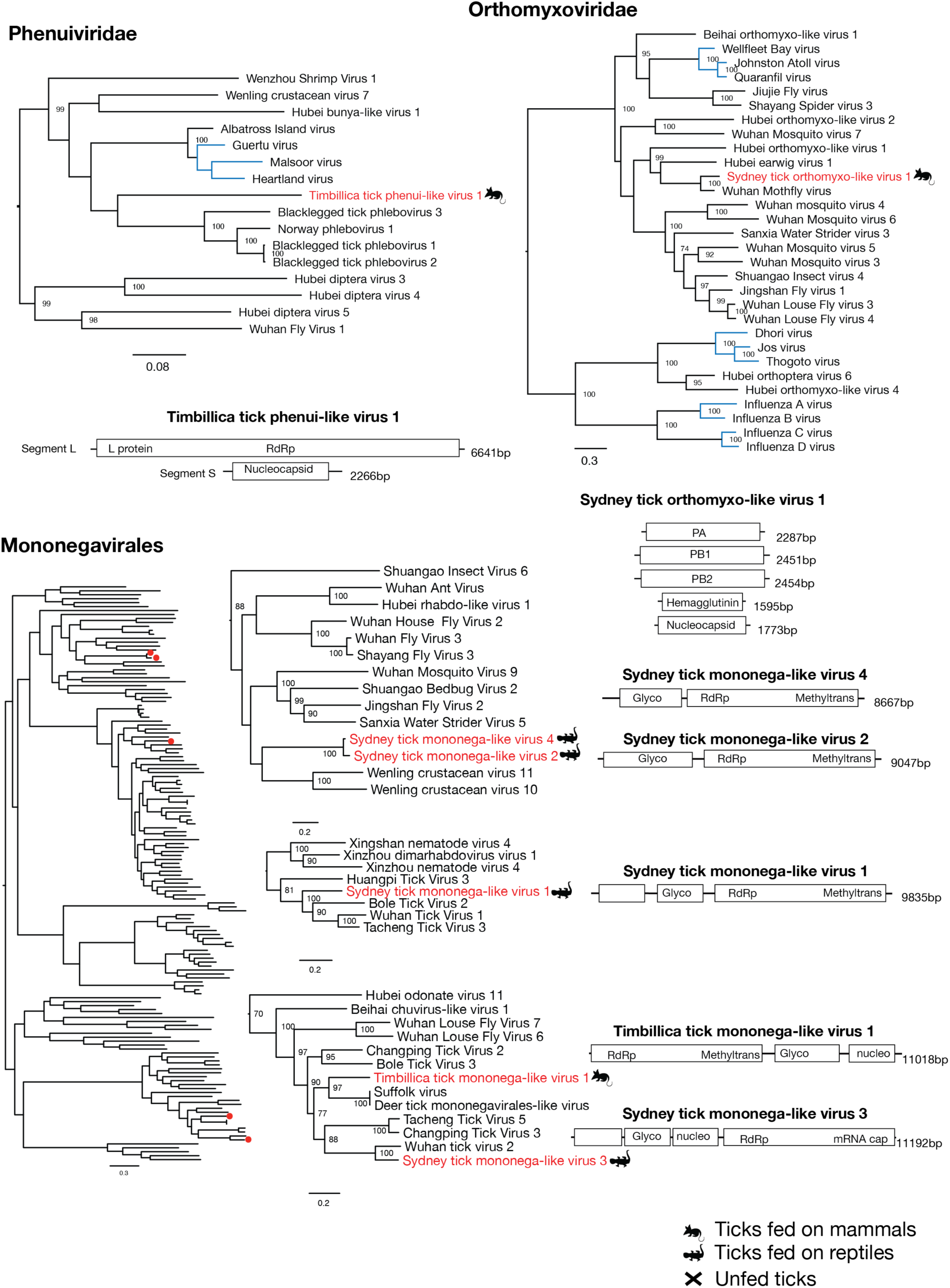
Phylogenetic relationships and genomic structure of the -ssRNA viruses sampled in this study. All trees are scaled according to the number of amino acid substitutions per site. The trees are midpoint rooted for clarity and bootstrap values (>70%) are shown. Viruses determined here are shown in red. Animal symbols denote the vertebrate host the tick was taken from or whether the virus was sampled from an unfed tick. Virus lineages previously shown to infect mammals or mammalian cells are shown in blue. Genome diagrams provide information on the length of each genomic segment, the number of predicted ORFs and predicted conserved protein structures.

The most well-represented order of viruses discovered in our survey were the *Mononegavirales*, as we identified five novel viruses clustering within the recently described *Chuviridae* family of arthropod-associated viruses (23). Timbillica tick Mononega-like virus 1 and Sydney tick mononega-like virus 3 clustered with a group containing Suffolk virus, Deer tick mononegavirales-like virus, and Wuhan tick virus 2 (Fig. 6). All of these viruses contained a single segment, with the number of ORFs varying from 3-4. Viruses belonging to the order *Mononegavirales* showed relatively high abundance in all the libraries in which they were identified. Sydney tick mononega-like virus 1 was closely related to other tick-associated viruses and represented 54% of the total viral reads in library 1 and 0.26% of total reads in the library. The genome is a single segment approximately 9835 nt in length that encodes three ORFs. Sydney tick mononega-like virus 4 and Sydney tick mononega-like virus 2 exhibited a close evolutionary relationship, with 95.7% amino acid identity between the two RdRp sequences. These viruses were highly divergent from those previously identified, and their sampled relatives are known to infect crustaceans (23). Sydney tick mononega-like virus 2 represented approximately 25% of total virus reads, while Sydney tick mononega-like virus 4 only represented approximately 1% of viral reads in this library. Both viral genomes encoded two ORFs. Timbillica tick mononega-like virus 1 and Sydney tick mononega-like virus 3 showed relatively close evolutionary relationships to previously described tick-associated viruses within the *Mononegavirales*, with Timbillica tick mononega-like virus 1 clustering with viruses associated with *Ixodes scapularis* and Sydney tick mononega-like virus 3 clustering with viruses identified in *Rhipicephalus microplus* (32, 33). Timbillica tick mononega-like virus 1 represented ~47% of total virus reads in the library in which it was identified, while Sydney tick mononega-like virus 3 represented approximately 18% of the viral reads identified in library 1. Timbillica tick Mononega-like virus 1 encoded three ORFs while Sydney tick mononega-like virus 3 encoded four (Fig. 6).

Finally, it is important to note that no reads matching the bacterium *B. burgdorferi* s.l. were identified in any of the BLAST searches of the RNA-Seq data generated here.

## DISCUSSION

We present the first metagenomic description of the viruses present in Australian ticks. Analysis of the transcriptomes of 146 ticks collected from two locations on the NSW coast revealed the capacity of these ticks to harbor a wide diversity of novel viruses. Previous meta-transcriptomic studies targeting arthropod viromes have revealed an abundance of novel and often highly divergent viruses (14, 23-25, 32). To date, however, there has been no study of the diversity of tick-associated viruses in Australia, nor of whether any of these viruses might play a role in tick-borne disease, despite the controversy surrounding tick-borne disease in this country (6, 7). Importantly, we also used the RNA-Seq meta-transcriptomic data generated here to search for the presence of known tick-associated pathogens, particularly the bacteria *B. burgdorferi* s.l. – the complex of bacterial species causing Lyme disease in parts of Europe and North America. Although we found evidence for a range novel bacterial species (data not shown), no *B. burgdorferi* s.l., was detected which is consistent with recent bacterial 16s rRNA-based metagenomic studies performed on hundreds of Australian ticks (11, 12). Hence, our results are in accord with the growing consensus that Lyme Disease is not present in Australia.

The abundance of the mitochondrial COX1 gene was not uniform across all libraries, although this is expected due to such factors as variation in the number of ticks included in each library, the size of each individual tick, and the extent to which the tick was engorged. It may also reflect the differing abundance of bacterial and fungal genetic material in the library. Importantly, however, the libraries were constructed based on the tick’s collection location, species, host, and the amount of RNA extracted from that tick, so that approximately equal amounts of RNA from each tick were included within a library.

Overall, we identified 19 novel viruses, one of which clearly merits additional investigation as a potential zoonotic pathogen based on its phylogenetic relationship to other tick-borne viruses associated with human disease. Specifically, Sydney tick colti-like virus 1 fell within the *Coltivirus* group within the family *Reoviridae*. This group currently comprises four virus species, two of which have been implicated in human tick-associated disease – Colorado tick fever virus and Eyach virus in North America and Europe, respectively (15, 16). Coltiviruses have also been isolated from rodents and bats, as well as the ticks that act as transmission vectors (30, 31). It is therefore possible that Sydney tick colti-like virus 1 is infecting the long-nosed bandicoots that acted as a distinct mammalian host, and may have the potential to cause zoonotic disease. This would obviously be of considerable concern given the close proximity of dense urban development to the national park in which these long-nosed bandicoots are found in Sydney. A systematic screening in long-nosed bandicoots and other mammalian species, as well as in human cases of tick-borne disease in Australia, is therefore warranted.

We also identified two novel viruses belonging to the *Flaviviridae* that clustered closely with rodent-associated viruses. This phylogenetic position, in combination with the extremely low abundance of these viruses, suggests that they were in fact derived from the tick’s vertebrate host, and were present in the blood contained in the engorged tick rather than being from the tick itself although this will need additional verification.

The majority of the ticks sampled and sequenced in this study were collected from National parks in the Sydney region, which is the most populated region in Australia. Encroachment of urban development on the habitats of native wildlife not only threatens the survival of native flora and fauna, but by changing the nature of the human-animal interface may also increase the risk of novel zoonotic disease. For example, changes in land use and encroachment of human development on bat habitats in sub-tropical areas of Australia have resulted in increased contact between fruit bats and horses, in turn leading to a spill-over of Hendra virus into the equine population (34). On a number of occasions, Hendra virus has been transmitted from horses to humans and resulted in fatal disease outcomes (35).

Of note was that Timbillica tick Phenui-like virus 1 groups with several other tick-associated viruses within the *Phenuiviridae*, and is related (albeit distantly) to Heartland virus that has been implicated in human disease in North America (36). Interestingly, another virus within this group, Albatross Island virus, was identified as a potential cause of a disease outbreak in the shy albatross (*Thalassarche cauta*) in north western Tasmania, Australia (21). The only previously described virus found within our data was the picornavirus *I holocyclus* iflavirus. This virus was previously identified in the salivary glands on *I. holocyclus* collected in the coastal regions of northern NSW and southern Queensland (28), and we detected it in the ticks collected from the Sydney region.

In sum, using a powerful meta-transcriptomics approach we identified 19 novel and sometimes divergent viruses within ticks collected in the central east coast of Australia, one of which is related to those already known to cause human disease. It is therefore clear that there is a great diversity of viruses in Australian wildlife that are yet to be discovered and as contact between urban areas and native wildlife increases, it is possible that some of these may pose a risk to public health and hence require careful monitoring.

## MATERIALS AND METHODS

### Ethics Statement

Ethics approval for this work was granted by the NSW Office of Environment and Heritage Animal Ethics Committee (number 000214/05) and from the Australian National University Animal Ethics Committee for the southern brown bandicoot sampled at the Timbillica state forests (A2015/26). Samples were collected under a scientific licence provided by the NSW Office of Environment and Heritage (number SL100104).

### Sample collection

Engorged ticks were collected live and immediately frozen in liquid nitrogen during population studies of bandicoots in Sydney’s North Head national park (33.8224° S, 151.2994° E) and Timbillica state forests (37.3712° S, 149.7211° E) south of Eden, NSW (Fig. 1). Specifically, ticks were collected from two marsupial species – long-nosed bandicoots (*Perameles nasuta*) in the north (Sydney) and southern brown bandicoots (*Isoodon obesulus*) in the south (Timbillica). As well as bandicoots, ticks were collected from the eastern blue-tongued lizard (*Tiliqua scincoides scincoides*) and both native (*Rattus fuscipes*) and invasive (*Rattus rattus*) rats unintentionally caught in treadle cage traps. Unfed *I. holocyclus* nymphs were collected at Katandra Bushland Sanctuary (33.6744° S, 151.2799° E) (Ingleside, Sydney) by flagging during five collection trips between May and September 2017. All the collected ticks were taken to the laboratory and identified using tick taxonomic keys (37).

### Extraction of RNA, pooling and sequencing

Ticks were washed in 1X Dulbecco’s phosphate-buffered saline (DPBS) solution and homogenised in 800μl lysis buffer using a TissueRuptor (Qiagen). Total RNA was then extracted using a RNeasy plus mini kit (Qiagen) according to manufacturer’s instructions. RNA quality was assessed using an Agilent 2100 bioanalyzer (Agilent Technologies). Samples were pooled for sequencing based on criteria including tick species, tick stage, sampling location and, where possible, the tick host species (Table 1). Host rRNA was first depleted using a Ribo-Zero-Gold rRNA removal kit (Human/Mouse/Rat, Illumina) and sequencing libraries were then prepared using a TruSeq total RNA library preparation kit (Illumina). Paired-end sequencing was performed on the Hiseq2500 platform (Illumina). Library preparation and sequencing was performed by the Australian Genomic Research Facility (AGRF).

### Assembly and analysis

Sequencing reads were trimmed for quality using trimmomatic and *de novo* assembled using Trinity (v.2.1.1) (38). The assembled contigs were subjected to BLAST searches against the NCBI nr protein database using diamond (v.0.9.10) (39) and contigs with significant sequence similarity to viral proteins were selected. Significant sequence similarity was determined by an e-value of less than 1e^-20^, contig length of more than 500 nucleotides, and an amino acid alignment length of more than 100 characters. Sequences were searched for predicted ORFs using the ExPASy Translate tool (https://web.expasy.org/translate/). The predicted ORF structure of the contig was then compared to the genome of the closest BLAST hit to determine if the contig was potentially an endogenous virus element rather than a true exogenous virus. The blast search results were also used to determine if any species of *Borrelia* bacteria, particularly *B. burgdorferi* s.l., were present in our data.

### Identification of host genes

The assembled contigs were subjected to blastn and blastx (v.2.6.0) (40) searches against the NCBI nt and nr databases and the output was used to identify the key marker gene cytochrome C oxidase subunit I (COX1) to confirm the species identification of the ticks in each library.

### Estimating transcript abundance

Ribosomal reads were identified by selecting all contigs showing nucleotide similarity of over 90% across a length of over 200 nucleotides and an e-value of less than 1e^-20^ to ribosomal reads within the NCBI nt database using a blastn (v.2.6.0) (40) search. Bowtie2 (v.2.2.5) (41) was then used to map sequence reads back to these contigs using the end-to-end alignment algorithm. Bowtie2 end-to-end alignment was used to map the sequence reads from each library to the viral sequences identified, as well as to the COX1 reference sequence from the host genome. The mapped alignments were then checked in Geneious (v.9.1.5) (42) to ensure the quality of the alignment and to find any potential mapping errors. The ribosomal read count was then subtracted from the total number of reads when calculating the abundance of viral and COX1 sequences.

### Phylogenetic analysis

To determine the evolutionary history of each virus, the RNA-dependent RNA polymerase (RdRp) or complete polyprotein of each virus was compared to previously described proteins from the relevant virus family. Accordingly, RdRp or polyprotein sequences representing the family of each virus discovered here were retrieved from NCBI RefSeq. The Mafft program (v.7.3.0.0) (43) was then used to align each group of sequences using the L-INS-i algorithm, with all ambiguously aligned regions removed using TrimAL (v.1.4.1) (44). This process resulted in a total of 10 sequence alignments of more than 380 amino acid positions each on which phylogenetic analysis could proceed. IQTree (v.1.6.1) (45) was used to determine the best-fit model of amino acid substitution based for each data set (which was found to be the LG model in all cases), and maximum likelihood phylogenetic trees were then estimated with the PhyML (v. 20150415) (46) program, using 100 bootstrapping replications to assess the support for each node.

### Accession numbers

All consensus virus sequences produced as part of this study have been submitted to the NCBI and assigned accession numbers XXXX-YYYY. All sequence reads generated in this study are available under the NCBI Short Read Archive (SRA) under accessions XXXX-YYYY.

## ACKNOWLEDGMENTS

This work is funded by an Australian Research Council (ARC) Australian Laureate Fellowship awarded to ECH (FL170100022). NL is funded by an ARC Future Fellowship (FT160100463), and EH is funded by an Australian Postgraduate Award. We thank all those who assisted with tick collection.

